# Comparative analysis of PknB inhibitors for reactivity and toxicity

**DOI:** 10.1101/423061

**Authors:** Nathan Wlodarchak, Jeff Beczkiewicz, Steven Seitz, Zhengqing Ye, John Feltenberger, J Muse Davis, Rob Striker

## Abstract

Bacterial serine/threonine kinases are increasingly sought after as drug targets for new antibiotics. PknB, an essential kinase in Mycobacteria tuberculosis, is intensely targeted, and many inhibitors are in the developmental pipeline. These inhibitors typically are derived from screens of known kinase inhibitors and most share similar chemical properties as their parent compounds were all designed for optimal pharmacokinetic properties in the human body. Here, we investigate the reactivity and toxicity of a proposed PknB inhibitor, YH-8, which does not follow traditional drug design rules. We found that the compound is highly reactive with thiolating agents and has appreciable toxicity in a zebrafish animal model. Furthermore, we find minimal anti-mycobacterial activity with non-tubercular mycobacteria strains. These data suggest that further investigation is needed into its efficacy and physiochemical properties if it is to be further developed as an effective antibiotic.

## INTRODUCTION

Drug development is a costly and time consuming endeavor which suffers from many roadblocks during the discovery process. Predicting which leads will be adaptable into clinical drugs and which hits are worth further investigation is a difficult task. Several sets of rules are commonly used to help with this task and reduce unnecessary investments into doomed compounds. Lipinski’s rules were developed to codify which physicochemical properties a compound should have to be effective at orally surviving in the human body and penetrating cells in order to be pharmacologically effective[1]. Several effective drugs do not follow these rules, but they have generally held true for the majority of clinically used oral drugs against human intracellular targets[2]. Although they can be a good guide, they certainly are not predictive of efficacy, and often hits are limiting so even those with poor Lipinski physicochemical properties but high efficacy can be worth further investigation and optimization. Similarly, rules identifying compounds that are highly reactive, promiscuous, or degraded are also widely applied to exclude compounds for further analysis. Pan-assay interference compounds (PAINS) were traditionally defined as compounds with reactive substructures that were commonly seen as hits across multiple biologic and biochemical assays[3]. PAINS reactivity can be easily calculated for known compounds, and scored Pass/Fail for a variety of drug-like filters (FAF drugs[4]) and used as a basis to remove reactive compounds from further biology experimentation. However, heavily relying on these rules may also exclude imperfect compounds that may still be useful[5], though it may be more challenging to advance these compounds forward than compounds without these undesirable reactivities.

The current antibiotic crisis has led to renewed interest in new anti-infectives, especially for emerging drug-resistant bacteria[6]. Unfortunately, commercial companies have few financial incentives for developing antibiotics, leaving much of the development to academic labs[7]. Academic labs generally do not have the resources that commercial companies possess, therefore, it is crucial to deploy smart drug development strategies to save time and money. While many academic labs are turning to computational methods for hit discovery and lead refinement; no single efficacious strategy has emerged as superior to traditional screening and selection processes. Still, computational analysis can be incredibly useful in identifying potential problems early in the drug discovery process. Additionally, high throughput assays to test reactivity and toxicity are useful in selecting hits that will have the most promising chance of becoming useful leads. Instead of using physicochemical property rules as dogma, a broader strategy may be to incorporate them into follow-up experiments.

Tuberculosis continues to be a significant public health threat with an estimated 9 million people developing tuberculosis in 2013, and 1.5 million deaths[8]. Inhibitors against new targets are needed to combat emerging resistant strains of Tb[9]. One of the most promising new targets is a eukaryotic-like serine/threonine kinase, PknB. PknB is a member of the Penicillin binding And Serine/Threonine Associated (PASTA) kinase family and is an essential gene product for mycobacteria[10]. PknB is important in detecting cell wall stress and remodeling and also is involved with metabolic homeostasis and host cell survival[11-14]. Several groups have discovered biochemical PknB inhibitors which have microbiologic activity against tuberculosis[15-18].

This study focuses on characterizing YH-8, a recently reported inhibitor for Tuberculosis[19]. While other PknB inhibitors reported adhere well to PAINS and Lipinski chemical properties, YH-8 revealed a characteristic highly reactive Michael acceptor, with recent organic synthesis methodology reports indicating its use as such[20-23]. Additional SAR studies by Xu *et al*. indicate that the double bond Michael acceptor linking the ketone and ester moieties was essential for its PknB inhibition. In our hands, the biochemical activity was far less potent than microbiologic activity[19], prompting further investigation. We discovered that it was not as potent against other mycobacteria as was reported for Mycobacterium tuberculosis, and the biochemical activity was highly dependent on assay conditions. We found that it adducted readily with a thiol reducing agent (DTT) and was toxic to zebrafish relative to another known non-specific kinase inhibitor. Collectively our data show that simple biochemical, reactivity, and toxicological tests on a promising hit compound can be used to identify downstream pharmacokinetic problems. Application of these tests will allow researchers to rapidly assess hit appropriateness for further development.

## RESULTS

### YH-8 Microbiologic activity

YH-8 (**Fig. 1**) was previously reported to inhibit *Mycobacterium tuberculosis* (Mtb) growth in broth with an MIC of 0.25 µg/mL[19]. We wanted to see whether we could reproduce similar results in other mycobacteria which are traditionally used for initial inhibitor screening. We tested this compound against *Mycobacterium smegmatis* and *Mycobacterium bovis*, Bacillus Calmette-Guerin (BCG) and were surprised to find both strains were more resistant than the reported MIC of Mtb. The MIC of YH-8 against M. *smegmatis* and BCG was 500 µg/mL and 100 µg/mL, respectively (**Fig. 2**). Although we were unable to test this compound against Mtb, it is highly surprising how poorly this compound performed against strains which are generally easier to inhibit growth relative to Mtb. Given this surprising result, we wanted to confirm our compound was supplied as requested. The certificate of analysis from chemical supplier Enamine as well as in-house NMR analysis found the purchased YH-8 conformed to the reported structure.

**Figure 1:**
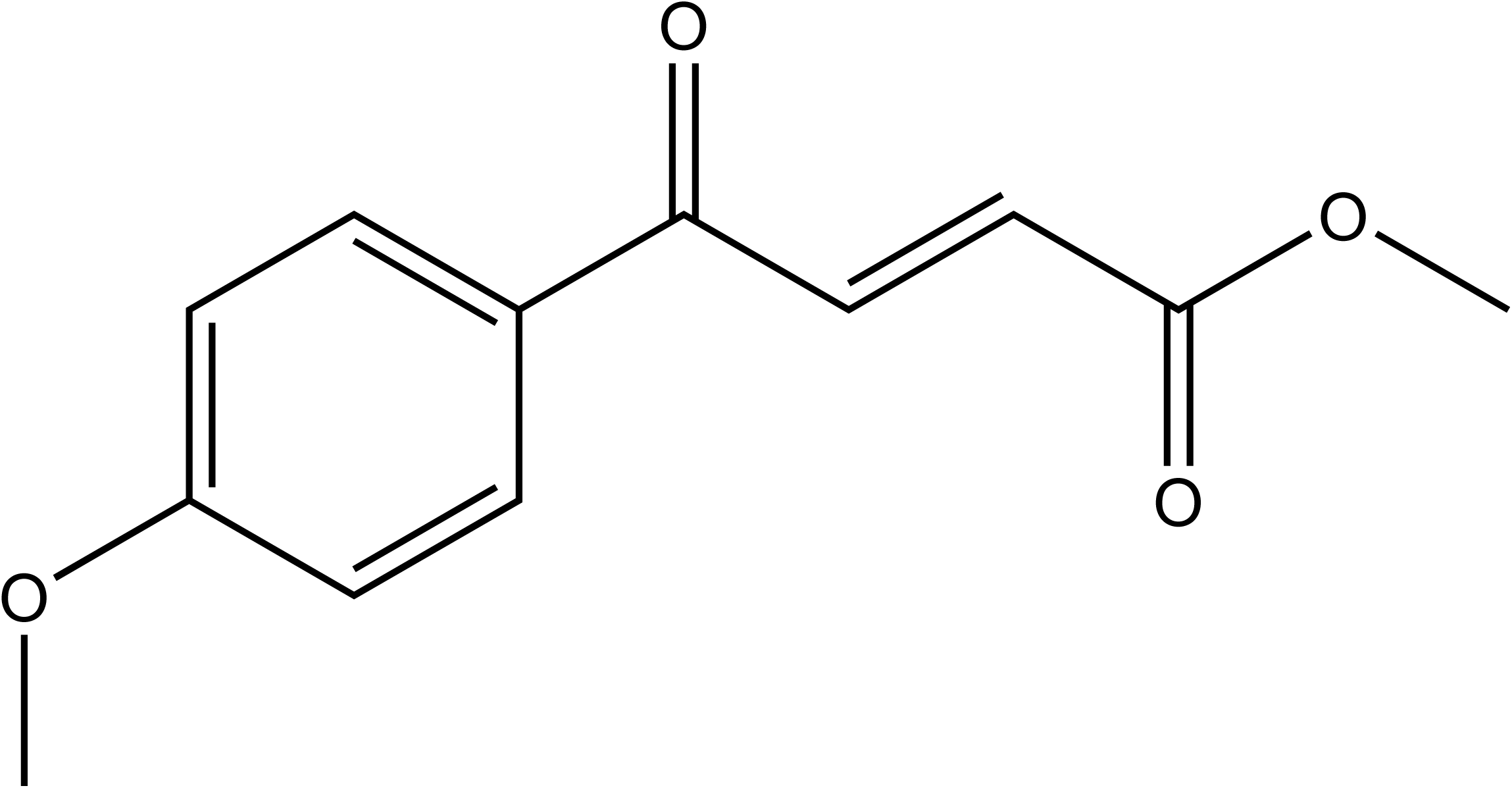
The structure of YH-8

**Figure 2:**
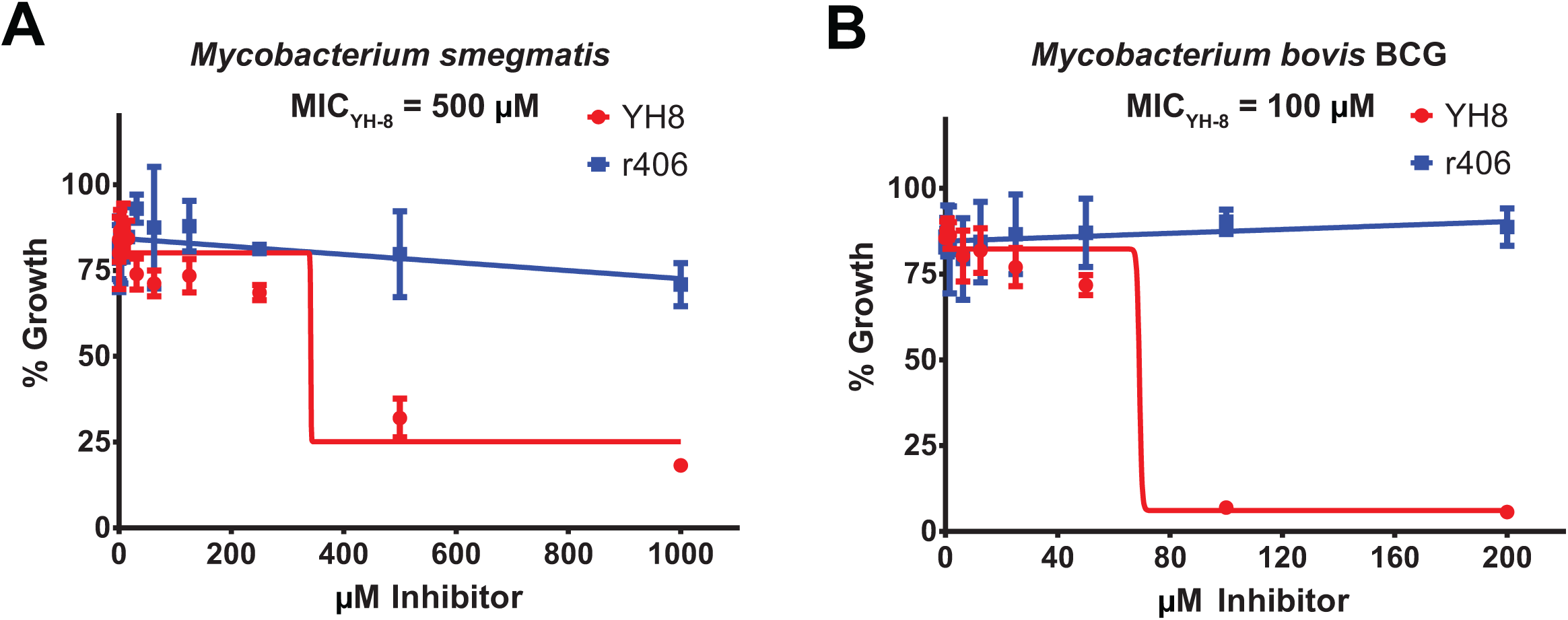
YH-8 has species specific inhibitory power. **A)** Mycobacterium smegmatis was treated at various doses of YH-8 and R406. R406 was unable to inhibit growth up to 1mM and YH-8 had a minimum inhibitory concentration (MIC) of 500 µM. **B)** Mycobacterium bovis (BCG) was more succeptable to YH-8 with an MIC of 100 µM.

### Biochemical inhibition of PknB by YH-8

PknB inhibition by YH-8 was previously reported at 20µM using GarA as a protein substrate[19]. We purified PknB and GarA as described in the methods and assayed the reaction inhibition by YH-8 compared to a non-specific S/T kinase inhibitor, R406[24]. We found the IC50 of YH-8 to be 6.6µM using our system (**Fig. 3A**). This difference may be largely due to differences between our purified PknB construct, as we used the entire cytoplasmic domain of the protein (1-331) as opposed to the structured kinase domain (1-279) as previously published[19]. This longer construct contains the juxtamembrane domain (280-331) which is required for full activation of the kinase[25]. Our typical assay buffer contains 1mM DTT to prevent oxidative damage to the proteins. When we carry out the reaction in the presence of 1mM DTT, all activity of YH-8 is lost (**Fig. 3A**). This result was also confirmed using a qualitative P32 kinase assay, which also visually confirms the substrate is phosphorylated in the presence of YH-8 and DTT (**Fig. 3B**). These data suggest that YH-8 has the ability to block kinase activity, but in a reducing environment, the adducted compound does not retain its activity.

**Figure 3:**
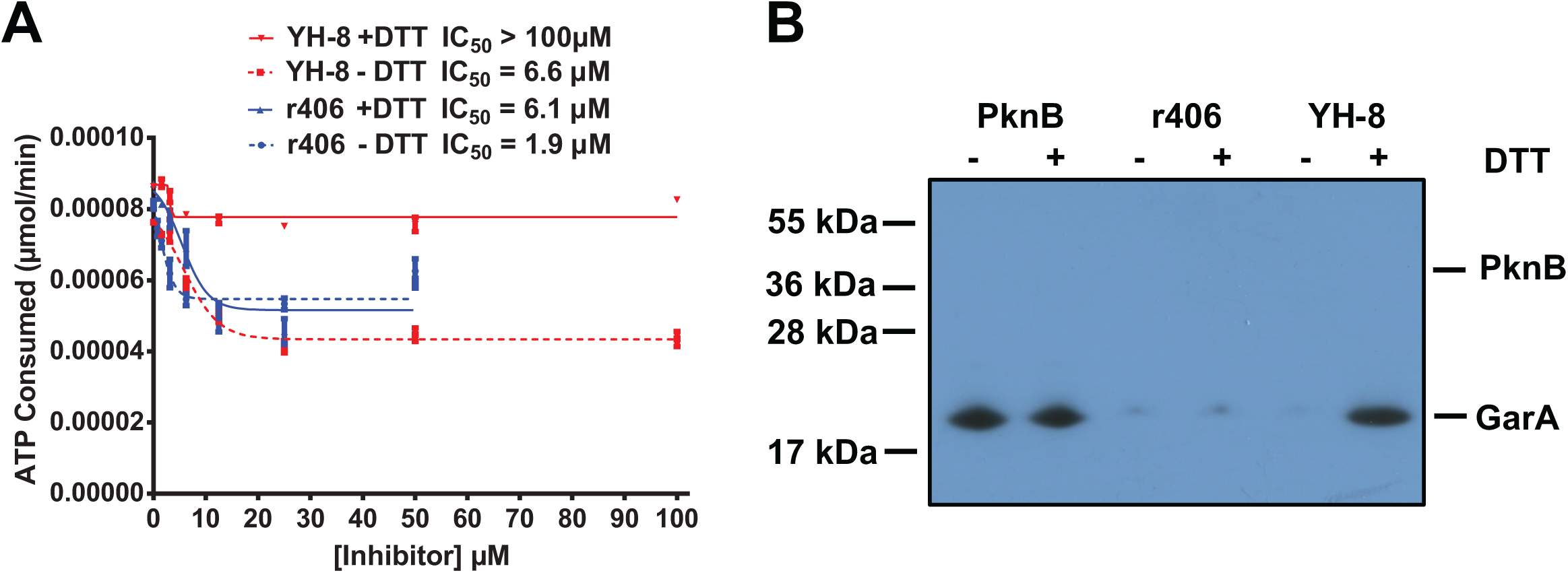
YH-8 cannot inhibit PknB in the presence of DTT. **A)** PknB enzymatic activity was assessed at various concentrations of YH-8 and a non-specific inhibitor, R406, using the ATP **Glo® assay in the presence (1mM) or absence of DTT.** YH-8 activity is completely abrogated in the presence of DTT. B) A radiometric phosphorylation assay visually confirms the results from A). YH-8 is unable to prevent GarA phosphorylation in the presence of DTT.

### YH-8 reactivity characterization

We noticed that the structure of YH-8 has a Michael acceptor (**Fig. 1**). We confirmed this by submitting the structure through a preprocessing drug filter (Free ADME-Tox Filtering Tool[4]) and found that the drug was a high risk Michael acceptor and would normally be rejected from any further screens. We wanted to assess YH-8 reactivity in a physiologically relevant chemical system. We used a DTT thiolation assay to determine its rate of reactivity and found that YH-8 was able to be rapidly thiolated in the presence of DTT and was able to make 1-2 adducts per molecule (**Fig. 4**). This is consistent with a highly reactive Michael acceptor and suggests the rate of inactivation for this compound would be very rapid in an *in vivo* system.

**Figure 4:**
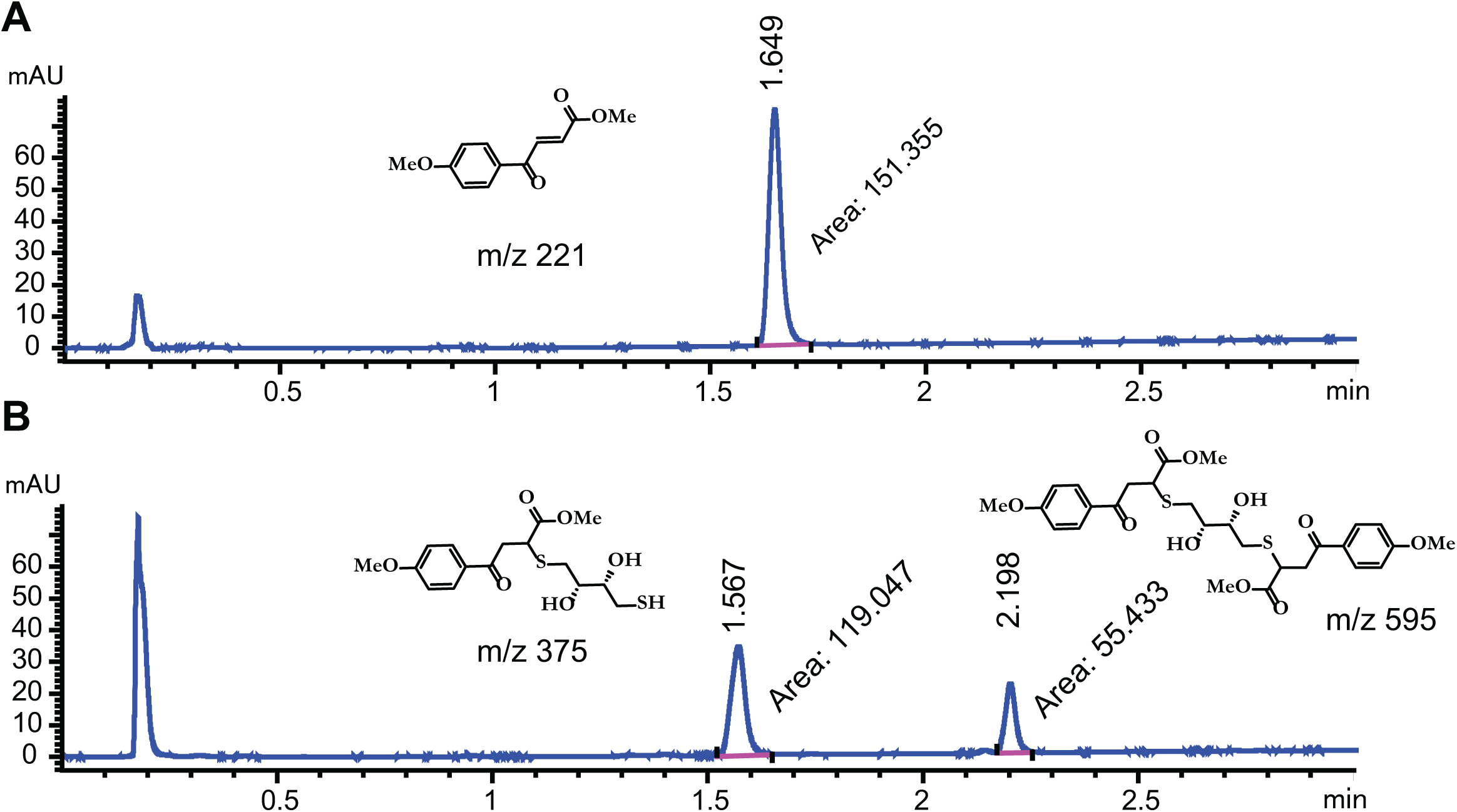
YH-8 is rapidly thiolated in the presence of DTT. YH-8 was assayed with DTT in an acetonitrile/PBS buffer as described in the methods. It was found that Michael addition of DTT to YH-8 was rapid: immediately after addition of DTT, an aliquot was diluted in 100% MeOH and injected on LCMS, where complete consumption of YH-8 was observed, and the single (M+H = 375) and double (M+H = 595) Michael products were observed.

### YH-8 toxicity in zebrafish

We hypothesized that this highly reactive compound would be toxic when used in model systems. Zebrafish (*Danio rerio*) are often used in the initial phase of drug toxicity testing due to their ease of use, cost, and ability to be assayed in a high throughput and rapid manner. Zebrafish are also excellent for bacterial kinase inhibitor testing since their kinome closely resembles that of humans and many kinases are activated during their development stages. We tested YH-8 and R406 for toxicity in zebrafish and found that YH-8 was highly toxic, killing every fish at every concentration we tested within the first 24 hours (**Fig. 5**). R406, despite its non-specificity, was considerably less toxic against zebrafish. This would suggest that YH-8 may be highly toxic at clinically relevant does needed for tuberculosis clearance and would likely fail clinical trials.

**Figure 5:**
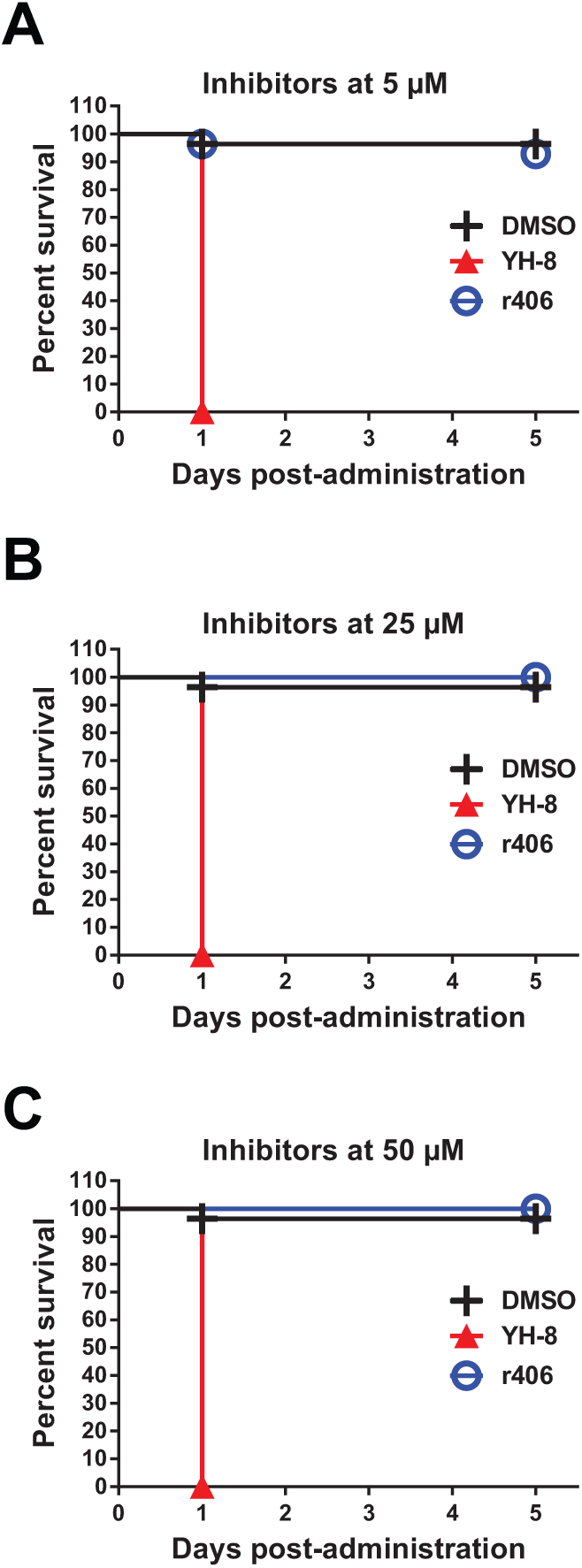
YH-8 is acutely toxic to zebrafish at biologically relevant doses. Zebrafish (Danio rerio) were exposed to **A)** 5 µM, **B)** 25 µM, or **C)** 50 µM of YH-8 or R406 for 5 days. At all doses tested, YH-8 was toxic to zebrafish within the first 24 hours. R406 did not have appreciable toxicity despite its non-specificity. DMSO was used as a control. Each experiment used nine animals and was done three times.

## DISCUSSION

Drug pharmacokinetics/pharmacodynamics are complex and problematic compounds are often not discovered until late in drug development. Several tools exist to pre-filter drug hits in order to eliminate some of the most problematic compounds before extensive time and money is invested in them. Certain structural features are conducive to false positive hits in many *in vitro* and in vivo screens, and collectively, these are referred to as PAINS[3]. Highly reactive compounds are not desirable due to the possibility of off target effects and toxicity. Although occasionally some of these compounds may not be as detrimental as predicted[5], the majority of the time they are a waste of resources. YH-8 would normally be discarded using pre-filtering databases for compound selection to eliminate a probably false positive. Our data clearly show the reactivity of YH-8 (**Fig. 4**) and differential activity under various redox environments (**Fig. 3**). It is also unknown whether YH-8 is actually competing with ATP or merely reacting with random residues on the kinase and reducing its activity. Curiously, previous reports show no inhibition of related mycobacterial kinases except for PknA, which would suggest that if the reactivity is non-specific, it is not complexing to other kinases in the same manner.

We observed considerably reduced microbiologic activity in other Mycobacteria (**Fig. 2**). This may be due to differences in inherent intracellular redox states and thiol compounds inherent to each species of mycobacteria, or the redox environment due to different growth conditions[26]. Furthermore, the human body contains numerous redox environments, including those rich with thiols such as glutathione[27], and as such, it is unlikely YH-8 would ever survive to reach its intended target. Curiously, previous reports demonstrated a lack of cytotoxicity with human macrophage cell lines and the ability to clear **Mycobacterium tuberculosis** infections from these cells[19]. Human macrophages are known to vary their content of glutathione and reactive oxygen species based on their activation and polarization. Glutathione itself also has bacteriostatic properties and is induced in the macrophage by tuberculosis infection. Altering glutathione levels may play a role in inactivating YH-8 and presence of YH-8 not affected by the glutathione flux may allow the balance to be tipped from a bacteriostatic state to a bactericidal state. Most importantly, not all cells have the protection from reactive species, such as YH-8, which is afforded to macrophages. This may explain the differences in toxicity between whole organisms, such as zebrafish (**Fig. 5**) and cell lines. Interestingly, the drug seems to be detectable in rat plasma, although the long term survival and morbidity of the rats was not commented on[28]. Although modifications may reduce this compound’s reactivity and toxicity, further evidence is needed to assess whether it will retain its effectiveness against *Mycobacterium tuberculosis.* Although YH-8 is reactive and has poor pharmacokinetic properties, several stable non-reactive inhibitors are known for PknB and other related bacterial kinases, including some which were in human trials for other diseases[15-17, 29-33]. Improving YH-8 by reducing its reactivity and toxicity and incorporating structure-activity relationship data from the aforementioned inhibitors may lead to improvements in the properties of YH-8, leading to a viable clinical lead compound.

## Materials and Methods

### Virtual analysis

YH-8 was built in ChemDraw (Perkin Elmer). The compound sdf file was submitted to the FAF-Drugs4 (Free ADME-Tox Filtering Tool)[4] server for analysis with default parameters. R406 was similarly built and analyzed as a control.

### Chemicals and reagents

All chemicals and reagents were sourced as previously published[29], with the following additions: R406 was obtained from Selleck Chemicals (Houston, TX) and YH-8 was obtained from Enamine (Kyiv, Ukraine). Tricaine was from Sigma (St. Louis, MO).

### Cell viability assays

#### Mycobacterium smegmatis

MC2155 WT *(Msmeg)* minimum inhibitory concentration (MIC) determinations were done according to a microdilution method. In a 96-well format, 50 µL of 1:1 serial diluted IMB-YH-8 or R406 was distributed to the plate in triplicate. The last column of the 96-well plate received medium without inhibitor. To correct for variations in DMSO concentration due to diluting, 40 µL of DMSO containing medium were added to the appropriate wells in specific concentrations to normalize all wells to 2% DMSO. Preparation of the bacterial culture to add was prepared beforehand in the following method. A frozen stock of *Msmeg* was streaked out onto Middlebrook 7H9 agarose plates supplemented with 10% ADC. After incubation at 37°C for two to four days, a colony was picked and inoculated into 2.5 mL of Middlebrook 7H9 broth supplemented with 10% ADC Middlebrook and 0.08% tyloxapol (v/v) and grown with shaking at 37°C. The next day, the culture was adjusted to an optical density of 0.3 (OD600), and 10 µL of the suspension was used to inoculate each well of microdilution plates. The plates were then incubated for 48 hours, stationary, at 37°C, and then 10 µL of 1mg/mL of resazurin dye was added. The plates were then incubated for an additional 12-15 hours at 37°C. Using a Synergy HT plate reader (Biotek) cell growth was quantified using fluorescence of the resazurin (excitation 530nm read 590nm). Measurements were scaled to the high wells that did not contain inhibitor and formatted as a percent relative to the high wells (100%).

#### Mycobacterium bovis

Bacillus Calmette-Guerin 1011 (BCG) MIC determinations were performed like those for *Msmeg* except for growth times. The 7H9 plates and broth cultures were incubated at 37°C for 4-5 days. When using the 96-well plates, the incubation time was 5 days at 37°C.

### NMR

^1^H NMR data was obtained from a Varian Inova 500 NMR spectrometer. The spectra were recorded in 10 mg cm-3 CDCl3 solutions with a probe temperature of ca. 300K and referenced to TMS. YH-8 **(*trans*-Methyl-4-(4-methoxyphneyl)-4-oxobut-2-enoate). 1H NMR** (CDCl3, 500 MHz) δ 8.06 (d, J = 9.0 Hz, 2H), 7.98 (d, J = 15.5 Hz, 1H), 7.10 (d, J = 9.0 Hz, 2H), 6.72 (d, J = 15.5 Hz, 1H), 3.87 (s, 3H), 3.78 (s, 3H)[34].

### Cloning & protein purification

Cloning and purification of PknB and GarA were done as previously published by our lab[29].

### Kinase assays

The kinase assays were performed as previously published[29] with the following changes: All reactions were done in 25µL volume. The buffer used for all kinase assays was 10mM Tris-HCl pH 7.4, 150mM NaCl and 1mM MgCl2 both with and without 1mM DTT. Drugs in 5mM DMSO were diluted in kinase buffer to 3/5 the final concentration using a serial dilution from 100 or 50µM to 0µM. The final DMSO concentration in the reactions was no more than 0.5%. Radiolabeled kinase assays were done as previously published[29] except the reaction buffer was the same as above (with and without DTT) and the inhibitors were assayed at 100 µM (YH-8) and 50 µM (R406).

### DTT thiolation assay

The DTT thiolation assay was performed on an Agilent 6120B LCMS equipment (Agilent PoroShell 120, EC-C18, 2.1 × 50 mm, 1.9 µM at 40 °C). LCMS studies were performed to monitor the reaction of YH-8 (100 μM) with DTT (500 μM) in 1: 1 CH3CN/PBS solvents. Samples were taken immediately after addition and diluted in 100% MeOH and injected on LCMS.

### Toxicity assays

Zebrafish were received from the breeding facility two days post fertilization (dpf) and dechorionated. Upon dechorionation, fish were transferred to fresh E3 zebrafish solution. Drugs were plated at 1.25x concentration to deliver 5, 25, or 50 μM final concentration in E3. Each drug concentration was repeated in nine wells for a total of 27 wells per drug per trial. Zebrafish were transferred with a micropipette using large orifice tips standardized at 20μL per well to give the final concentrations of drugs with fish at 1x. The total volume of fluid is 100μL in each well. Outermost wells were filled with DI water to limit evaporation in test wells. Embryos were incubated at 28.0°C for five days. Plates were observed every 24 hours and presence of a regular heartbeat was considered the threshold for survival. Fish alive at the completion of the experiment were euthanized by placing fish in a bath of E3:bleach at a 5:1 ratio. The bath was then incubated at room temperature for five minutes before the solution was discarded per our animal use protocol.

## Acknowledgments

We would like to thank Adel Talaat for providing us with the M. smegmatis and BCG strains, and Becky Procknow and Yen Tuong for help with cloning and protein purification. We would particularly like to thank Andrew Mehle for allowing us to use his ÄKTApurifier.

This work was supported by grants from the Hartwell Foundation and the Department of Veterans Affairs Merit award. This study made use of instrumentation at the Medicinal Chemistry Center at UW-Madison funded by the UW School of Pharmacy.

## Author Contributions

N.W. oversaw the project, designed the experiments, performed biochemistry experiments, analyzed the data, rendered the figures, and wrote the paper. J.B. collected and analyzed *M. smegmatis* and BCG data. S.S. performed zebrafish experiments under the direction of M.D. Z.Y. performed NMR and chemical stability assays under the direction of J.F. R.S. conceived and oversaw the project, edited the manuscript, and provided direction for writing and submission.

